# Facilitation of sensorimotor temporal recalibration mechanisms by cerebellar tDCS in patients with schizophrenia spectrum disorder and healthy subjects

**DOI:** 10.1101/2023.09.28.559952

**Authors:** Christina V. Schmitter, Benjamin Straube

## Abstract

Core symptoms in patients with schizophrenia spectrum disorder (SSD), such as hallucinations or ego-disturbances, have been associated with a failure of the forward model to adequately predict the sensory outcomes of self-generated actions. Importantly, depending on the requirements of the environment, forward model predictions must also be able to recalibrate flexibly, for example to account for additional delays between action and outcome. In this study, we aimed to investigate whether non-invasive brain stimulation via transcranial direct current stimulation (tDCS) can be used to improve these sensorimotor temporal recalibration mechanisms in patients and in healthy subjects.

While receiving tDCS on the cerebellum, temporo-parietal junction (TPJ), supplementary motor area (SMA), or sham stimulation, patients with SSD and healthy control subjects were repeatedly exposed to delays between actively elicited or passively performed button press movements and auditory sensory outcomes. Effects of this procedure on temporal perception were assessed with a delay detection task.

We found similar sensorimotor temporal recalibration effects in both SSD and healthy subjects. Furthermore, cerebellar tDCS facilitated recalibration effects in both groups.

Our findings indicate that sensorimotor recalibration mechanisms may be preserved in SSD and highlight the importance of the cerebellum in both patients and healthy subjects for this process. Our results suggest that cerebellar tDCS could be a promising tool for addressing deficits in action-outcome monitoring and related adaptive sensorimotor processes in SSD, and potentially alleviating associated symptoms.

## 1 Introduction

Core symptoms in patients with schizophrenia and schizoaffective disorder (referred to as schizophrenia spectrum disorders, SSD) encompass hallucinations (e.g., perceiving the own inner speech as external voice) and ego-disturbances (e.g., perceiving own thoughts or actions as externally controlled). The emergence of these symptoms has been associated with a failure of adequately predicting the sensory outcomes of one’s own actions.^1^ When performing an action, copies of the motor commands are thought to be used by an internal forward model to predict the action’s sensory outcomes. Re-afferent sensory input that is in line with the prediction is typically associated with modulations in neural responses in multiple brain regions compared to input that deviates from the prediction, and is thus perceived as self-generated.^2–4^ Dysfunctions in this predictive mechanism can result in the misattribution of self-generated sensory input as externally produced and lead to the symptoms in SSD described above.^5–10^

Importantly though, forward model predictions are not rigid, but need to be able to flexibly recalibrate to preserve adequate distinction between self- and externally generated input under dynamically changing environmental conditions. For instance, the outcome of an action can be transiently delayed under certain circumstances, e.g., a mouse click can lead to delayed responses from a computer.^11^ Studies with healthy subjects have shown that after repeated exposure to a delayed action-outcome, the predicted outcome timing shifted toward that delay. As consequence, the delayed outcome was perceived as occurring in synchrony with the action, (known as sensorimotor temporal recalibration effect, TRE), and neural responses for the delayed outcome resembled the ones typically observed for undelayed outcomes.^17^ To date, it remains unknown whether the dysfunctions in predictive mechanisms in SSD are due to a general failure of prediction generation or, more specifically, a failure of adequately recalibrating these predictions to the changes in action-outcome delays.

Neural correlates of the predictive processes based on the forward model have been identified in several brain regions. The cerebellum is most prominent in this regard since it has been suggested to play a vital role in generating and updating predictions about sensory action-outcomes.^2,4,18–22^ Additionally, regions in parietal cortex, particularly the temporo-parietal junction (TPJ) or angular gyrus,^9,23–25^ and the supplementary motor area (SMA)^26,27^ could be associated with the subjective feeling of control or agency over action-outcomes and the distinction between self- and externally generated stimuli. Interestingly, non-invasive brain stimulation techniques have demonstrated the potential to modulate these processes. For instance, transcranial magnetic stimulation (TMS)^28^ or transcranial direct current stimulation (tDCS)^29^ of the cerebellum influenced the effect of sensorimotor temporal recalibration on perception in healthy subjects. Furthermore, tDCS on the angular gyrus,^30^ the pre-supplementary motor area,^31,32^ and frontal regions^33,34^ modulated measures for agency and action-outcome-related processing, even in SSD.^35^ Thus, tDCS may be a promising tool to enhance sensorimotor temporal recalibration mechanisms and thereby improve action-outcome processing and self-other distinction in SSD.

Therefore, the present study investigated whether sensorimotor temporal recalibration mechanisms are impaired in SSD compared to healthy control (HC) subjects, and whether tDCS on the bilateral cerebellum, right SMA, or right TPJ can enhance recalibration and thus reduce potential deficits. Subjects were exposed to delayed or undelayed tones following either actively or passively elicited button press movements, and the effects of this procedure on auditory and visual temporal perception were assessed with a delay detection task. While active movements were expected to trigger sensorimotor temporal recalibration based in the forward model, passive movements controlled for recalibration effects due to changes in the expected inter-sensory timing between the tactile sensations during the button movement and the auditory or visual outcome.^36–39^ We expected patients to exhibit reduced temporal recalibration compared to HC, specifically in active conditions, due to impaired recalibration of forward model predictions. We expected tDCS applied on the mentioned brain regions to enhance temporal recalibration in both groups, particularly in active conditions, due to the presumed importance of these regions in generating forward model predictions.

## 2 Materials and methods

### 2.1 Participants

Twenty-four patients with SSD and 20 HC with no psychiatric diagnosis (10 female, mean age: 36.90, *SD* = 10.37) participated in the study. Two patients had to be excluded (see supplementary material S1), resulting in a final sample of 22 patients (11 female, mean age: 35.80, *SD* = 10.37). Fifteen patients were diagnosed with an ICD-10 diagnosis of schizophrenia, six patients with schizoaffective disorder, and one patient with an acute and transient psychotic disorder (for further details on sample characteristics see supplementary material S1). All subjects had normal or corrected-to-normal visual acuity, normal hearing, and no history of neurological disorders. No contraindications for tDCS (e.g., electric, or metallic implants) were reported. Subjects gave written informed consent and were financially reimbursed for their participation. The study was conducted according to the Declaration of Helsinki and was approved by the local ethics commission (Study 06/19) of the medical faculty of University of Marburg, Germany. The study was pre-registered in the German Clinical Trials Register (DRKS-ID: DRKS00025885).

### 2.2 Transcranial direct current stimulation

tDCS was applied using a DC stimulator (neuroConn GmbH, Ilmenau) and two rubber electrodes (5 x 7cm) in saline-soaked sponges (0.9% NaCl). For all stimulation conditions, the anode was placed over the respective brain region since anodal tDCS has been shown to increase cortical excitability.^40^ The cathode was attached on the deltoid muscle of the right upper arm (see supplementary material S2 for details on the electrode placement). All electrodes were attached with rubber bands. The stimulation was applied with a current of 2mA for 20 minutes (+ 10 seconds fade in and fade out periods). Next to these three active stimulation conditions, a sham stimulation condition was implemented by using sinus (HW) mode for 30 seconds. The stimulation parameters were chosen in accordance with established tDCS safety guidelines.^41^ Each subject experienced the four stimulation conditions (cerebellum, SMA, TPJ, sham) in four separate sessions. Sessions were performed at least 18 hours apart to prevent residual effects from the previous stimulation. The stimulation conditions were applied in counterbalanced order, ensuring that in each group, across subjects, each of the four stimulation conditions was applied approximately equally often during the first, second, third, or fourth session. Subjects were sequentially assigned to one of the possible combinations of stimulation conditions and were unaware of the hypothesized effects of stimulating the respective brain region on task performance.

### 2.3 Equipment and stimuli

Subjects performed the experiment in a dimly lit room in front of a computer screen. Button presses were executed with the right index finger using a custom-made electromagnetic passive button device. In active conditions, subjects pressed the button actively by themselves. In passive conditions, the button was pulled down automatically by an electromagnet (max. force 4N). An elastic fabric band was used to attach the subjects’ fingers to the button to ensure that it smoothly followed its movement in passive conditions. When the button reached the lowest position, the presentation of an auditory or visual stimulus was triggered. The visual stimulus was a Gabor patch (1° visual angle, spatial frequency: 2 cycles/degree) which was presented at the center of the screen. The auditory stimulus was a brief sine-wave tone (2000Hz with 2ms rise and fall) presented through headphones. Both stimuli appeared for a duration of 33.4ms. All stimuli were created and presented using Octave and the Psychophysics Toolbox.^42^ To prevent any influence of the direct visual or auditory feedback from the actual button presses on sensory outcome perception, the button device was covered by a black box and pink noise was applied through headphones during the experiment.

### 2.4 Experimental design and task description

The experiment consisted of multiple pairs of adaptation and test phases. In adaptation phases 18 consecutive button presses had to be executed each followed by the tone as auditory sensory outcome. The button presses were either performed actively or they were elicited passively (factor *movement type*). The tone occurred either immediately after the button press (undelayed, 0ms delay) or was delayed by 200ms (factor *adaptation delay*). The undelayed tones should align with the natural prediction of undelayed action-outcomes, while exposure to the delayed tones was expected to induce the need for temporal recalibration. Originally, we chose an adaptation delay of 150ms as is has been used in previous studies with young healthy subjects.^29,43^ But due to the generally lower delay detection performance of the on average older subjects in this study, we increased the adaptation delay to 200ms after inspecting the data of the first four patients. Nonetheless, the data of all patients were included in the current analyses. Pairwise comparisons indicated that excluding the data of these four patients would not lead to differences in group-dependent effects in overall temporal recalibration.

Each adaptation phase was followed by a test phase that assessed the impact of the adaptation delay on perception. A test phase consisted of six test trials for which the button had to be pressed once, either actively or passively. The movement type was the same as the one used in the previous adaptation phase. While the stimulus during adaptation phases was always auditory, in test phases, the button presses elicited either the auditory or the visual stimulus (factor *test modality*). This was done because the TRE has previously been shown to transfer between modalities, such that recalibration to a sensorimotor delay in one modality also affected temporal perception in another modality.^13,43–45^ The visual or auditory stimulus occurred with one of six test delay levels (0, 83, 167, 250, 333, 417ms). Each of the test delays was used once in each test phase in counterbalanced order. Subjects were instructed to report via keyboard presses after each trial whether they detected a delay between the button press and the stimulus. The TRE was defined as the difference in the proportion of detected delays after exposure to delayed vs. undelayed tones with worse performance for delayed tones reflecting a shift of the expected stimulus timing in the direction of the adapted delay, indicating temporal recalibration.

### 2.5 Procedure

An adaptation phase started with instructions displayed for 2000ms indicating the movement type of the button presses (see **Fig. 1**). After the instructions disappeared, subjects could start pressing the button or it started to move passively. Each button press was followed by the tone, either undelayed or delayed by 200ms. After nine button presses, a fixation cross appeared on the screen for a jittered duration (1000, 1500, 2000, or 2500ms) indicating a short break. After the fixation cross disappeared, the remaining nine button presses could be performed.

**Fig. 1.**
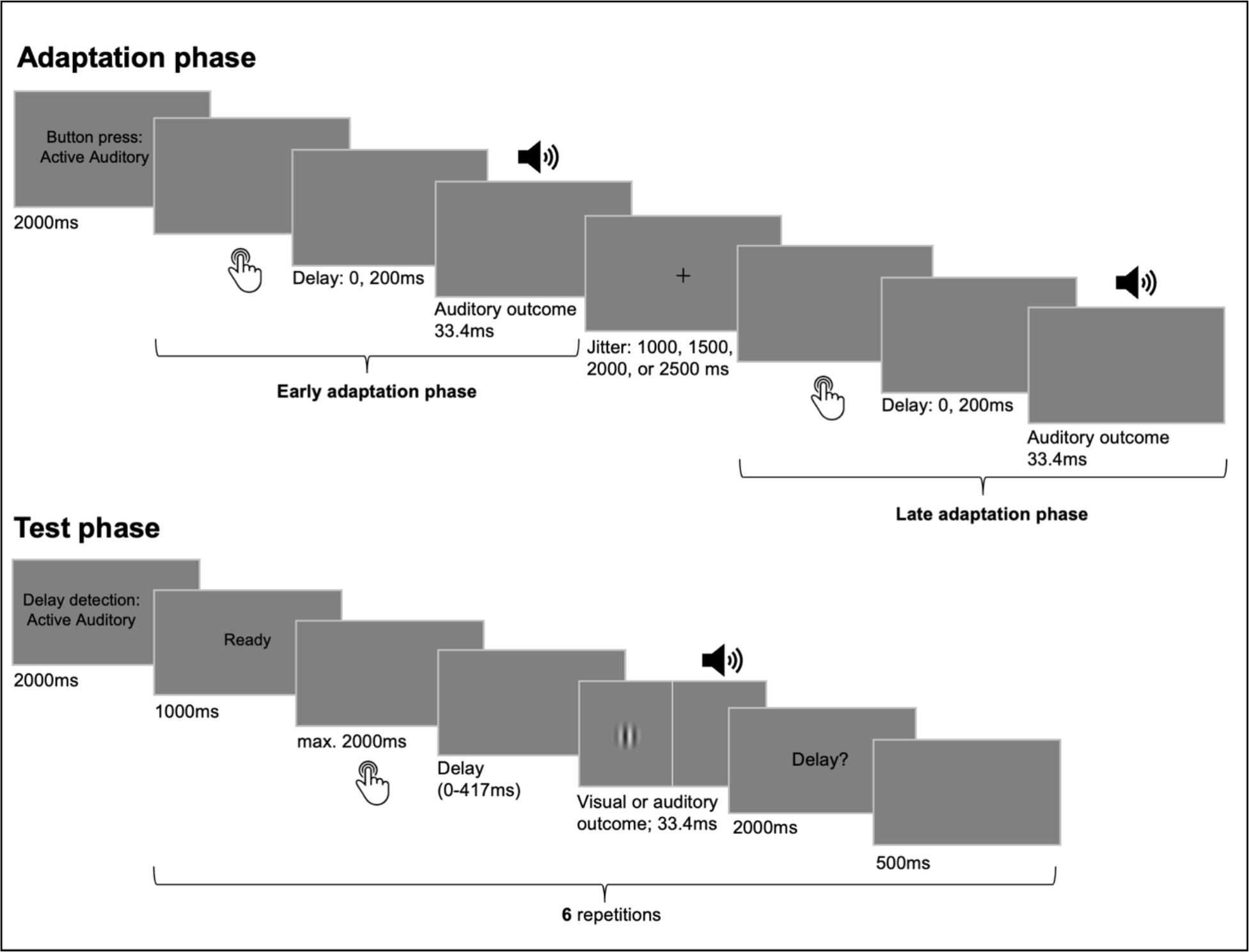
Trial sequence and timing of events. The experiment consisted of multiple pairs of adaptation and test phases. During adaptation phases, 18 button presses had to be performed either actively by the subjects or they were executed passively. A button press was followed by a delayed (200ms) or undelayed tone. Adaptation phases were divided into two parts separated by a fixation cross presentation. In test phases, the button was pressed once in each test trial, either actively or passively. Here, the outcome occurred after one of six delays (0-417ms) and subjects had to report in each trial whether they detected a delay. The outcome modality was always auditory during adaptation, but it could be visual or auditory during test.

A test phase was initiated by instructions (2000ms) about the movement type and stimulus modality of the following test trials. The cue “Ready” was presented for 1000ms before each test trial. After the cue disappeared, subjects had 2000ms in active conditions to press the button. But they were instructed to wait for approx. another 700ms to ensure that the movement was voluntary and not reflexive upon cue disappearance.^35,46^ Passive movements were initiated after a jittered interval of 0, 500, 1000ms. Each button press triggered the auditory or visual outcome with one of the six test delays. Afterwards, the question “Delay?” was presented and subjects had 2000ms to respond via keyboard press whether they detected a delay. After a pause of 500ms the “Ready” cue initiated the next test trial. After the last trial of a test phase a jittered inter-trial interval (1000, 1500, 2000, 2500ms) was inserted before the start of the next adaptation-test pair.

Each of the eight experimental conditions was presented with eight adaptation-test pairs per session. Within each session, conditions with the same adaptation delay were blocked to prevent spill-over effects due to rapid switching of delays. The first block of conditions with one of the adaptation delays took place while the stimulation was applied. After this first task block, electrodes were detached. The conditions with the other adaptation delay were presented in a second task block without stimulation. Whether the undelayed or delayed tones were presented first as well as the order of conditions within blocks was counterbalanced across subjects.

After each session, side effects due to the tDCS stimulation (e.g., itching sensations, headache, changes in visual perception, difficulties in concentration) had to be rated on a scale from one (no side effect) to five (strong side effect) using a custom-designed questionnaire of 28 items. During the first session, subjects additionally went through a training procedure to familiarize with the task (see supplementary material S3).

### 2.6 Data analyses

Test trials for which no button press (SSD: 1.858%, HC: .781% of all trials) or response (SSD: 5.617%, HC: 3.503% of all trials) was registered were excluded from the analyses. The percentage of detected delays served as a measure for delay detection performances and was calculated for each subject and experimental condition. These data were forwarded into a mixed ANOVA with the between-subjects factor *group* and the within-subjects factors *stimulation, test modality, movement type*, and *adaptation delay*. We examined main and interaction effects including the factors *adaptation delay* and *group*, to test for the impact of the adaptation delay on temporal perception and the modulatory influence of the other experimental factors as well as group-dependent effects. For significant interaction effects with the *adaptation delay*, we calculated the TRE defined as the difference in the percentage of detected delays in conditions with the 200ms compared to 0ms adaptation delay. Positive values indicate worse detection performance after exposure to the 200ms delay, reflecting a TRE into the expected direction. If indicated, post-hoc one-sided one-sample t-tests were used to assess whether the TRE was significantly greater than zero in the individual conditions of an interaction effect. Furthermore, two-samples t-tests were used to specify the difference in the TRE between the relevant conditions or groups. Since we had clear hypotheses regarding the direction of difference in TRE between conditions (i.e., a stronger TRE in HC vs. SSD, active vs. passive, auditory vs. visual conditions, and in active tDCS conditions vs. sham stimulation) one-sided t-tests were used. All post-hoc tests were Bonferroni-corrected if indicated. For significant interaction effects without the group factor, post-hoc tests were exploratorily performed not only across but also individually for both groups to identify similarities in temporal recalibration effects for SSD and HC.

Finally, we tested for differences in perceived stimulation side effects between the groups and stimulation conditions by a mixed ANOVA with the factors *group* and *stimulation* (results are reported the supplementary material S5). All analyses were performed in JASP (Version 0.14.1).^47^

## 3 Results

The ANOVA on the percentage of detected delays revealed no significant main effect of *group* [*F*(1, 40) = .239, *p* = .628, η_p_^2^ = .006], indicating that patients did not differ from HC in general delay detection abilities [HC: *Mean* = 37.138, *SD* = 8.845; SSD: *Mean* = 41.758, *SD* = 17.393]. Furthermore, none of the interaction effects including the factors *group* and *adaptation delay* were significant (all *p* > .139), thus providing no evidence for impairments in temporal recalibration and differences in the effectiveness of tDCS in patients with SSD* (see supplementary material S4 for a summary of all effects).

However, across groups and conditions, there was a significant main effect of the *adaptation delay* [*F*(1, 40) = 14.033, *p* < .001, η_p_^2^ = .260]. Thus, subjects’ perception recalibrated to the 200ms delay between button press and auditory outcome, leading to a significant TRE in terms of reduced delay detection performance [*Mean TRE* = 3.197, *SD* = 5.439; see **Table 1** for an overview of effects computed individually for both groups]. Additionally, the interaction of *movement type* and *adaptation delay* was significant [*F*(1, 40) = 8.762, *p* = .005, η_p_^2^ = .180]. Post-hoc tests revealed that the TRE was significantly greater than zero in both active [*Mean TRE* = 4.103, *SD* = 4.586, *t*(41) = 5.799, *p* < .001, *d* = .895, ɑ_corr_ = .025] and passive conditions [*Mean TRE* = 2.291, *SD* = 6.830, *t*(41) = 2.174, *p* = .018, *d* = .335, ɑ_corr_ = .025], but was significantly stronger in active ones [*Mean difference* = 1.812, *SD* = 4.124, *t*(41) = 2.848, *p* = .003, *d* = .439]. Furthermore, there was a significant *test modality* and *adaptation delay* interaction [*F*(1, 40) = 9.229, *p* = .004, η_p_^2^ = .187]. While the TRE was significantly greater than zero for both, audition [*Mean TRE* = 4.279, *SD* = 5.939, *t*(41) = 4.669, *p* < .001, *d* = .720, ɑ_corr_ = .025] and vision [*Mean TRE* = 2.115, *SD* = 5.871, *t*(41) = 2.335, *p* = .012, *d* = .360, ɑ_corr_ = .025], indicating a modality transfer of the TRE, it remained significantly larger in auditory (unimodal) than in visual (cross-modal) conditions [*Mean difference* = 2.164, *SD* = 4.599, *t*(41) = 3.050, *p* = .002, *d* = .471]. The interaction of the three factors *movement type, test modality*, and *adaptation delay* [*F*(1, 40) = 7.781, *p* = .008, η_p_^2^ = .163] further indicated that the active-passive difference in the TRE was specific to auditory outcomes [*Mean difference* = 4.187, *SD* = 5.872, *t*(41) = 4.622, *p* < .001, *d* = .713, ɑ_corr_ = .025; Active: *Mean TRE* = 6.373, *SD* = 6.255, *t*(41) = 6.603, *p* < .001, *d* = 1.019, ɑ_corr_ = .025; Passive: *Mean TRE* = 2.185, *SD* = 6.976, *t*(41) = 2.030, *p* = .024, *d* = .313, ɑ_corr_ = .025] but did not transfer to the visual modality [*Mean difference* = -.563, *SD* = 7.625, *t*(41) = -.478, *p* = .682, *d* = -.074, ɑ_corr_ = .025; Active: *Mean TRE* = 1.834, *SD* = 4.662, *t*(41) = 2.549, *p* = .007, *d* = .393, ɑ_corr_ = .025; Passive: *Mean TRE* = 2.396, *SD* = 8.733, *t*(41) = 1.778, *p* = .041, *d* = .274, ɑ_corr_ = .025].

**Table 1.** TREs for individual conditions and comparisons of conditions, evaluated individually for both groups.

| TRE | Group | Mean+/-SD | t-value | p-value | $\alpha_{\text{corr}}$ | Cohen's d |
| --- | --- | --- | --- | --- | --- | --- |
| Across conditions | HC | **2.218 +/- 4.033** | **2.459** | **.011\*** | **.05** | **.532** |
|  | SSD | **4.087 +/- 7.684** | **2.495** | **.012\*** | **.05** | **.550** |
| Active | HC | **3.584 +/- 3.233** | **4.959** | **< .001\*\*\*** | **.025** | **1.109** |
|  | SSD | **4.575 +/- 7.826** | **2.742** | **.006\*\*\*** | **.025** | **.585** |
| Passive | HC | .852 +/- 5.802 | .656 | .260 | .025 | .147 |
|  | SSD | 3.599 +/- 8.733 | 1.933 | .033 | .025 | .412 |
| Active > Passive | HC | **2.733 +/- 4.813** | **2.539** | **.010\*** | **.05** | **.568** |
|  | SSD | .976 +/- 6.231 | .735 | .235 | .05 | .157 |
| Auditory | HC | **3.883 +/- 4.283** | **4.055** | **< .001\*\*\*** | **.025** | **.907** |
|  | SSD | **4.639 +/- 8.528** | **2.551** | **.009\*\*** | **.025** | **.544** |
| Visual | HC | .553 +/- 4.878 | .507 | .309 | .025 | .113 |
|  | SSD | 3.535 +/- 8.196 | 2.023 | .028 | .025 | .431 |
| Auditory > Visual | HC | **3.331 +/- 4.382** | **3.399** | **.003\*\*** | **.05** | **.760** |
|  | SSD | 1.103 +/- 6.605 | .784 | .442 | .05 | .167 |
| Auditory: Active | HC | **5.816 +/- 3.905** | **6.661** | **< .001\*\*\*** | **.025** | **1.490** |
|  | SSD | **6.879 +/- 10.410** | **3.099** | **.003\*\*** | **.025** | **.661** |
| Auditory: Passive | HC | 1.950 +/- 6.342 | 1.375 | .092 | .025 | .308 |
|  | SSD | 2.399 +/- 9.244 | 1.217 | .119 | .025 | .259 |
| Visual: Active | HC | 1.353 +/- 4.391 | 1.278 | .092 | .025 | .308 |
|  | SSD | 2.271 +/- 7.292 | 1.461 | .079 | .025 | .311 |
| Visual: Passive | HC | -.247 +/- 7.377 | -.150 | .559 | .025 | -.033 |
|  | SSD | 4.799 +/- 10.894 | 2.066 | .026 | .025 | .441 |
| Auditory:<br>Active > Passive | HC | **3.866 +/- 6.128** | **2.821** | **.005\*\*** | **.025** | **.631** |
|  | SSD | **4.480 +/- 9.834** | **2.137** | **.022\*** | **.025** | **.456** |
| Visual:<br>Active > Passive | HC | 1.600 +/- 7.226 | .990 | .167 | .025 | .221 |
|  | SSD | -2.528 +/- 8.659 | -1.370 | .907 | .025 | -.292 |
| Sham | HC | 1.160 +/- 5.923 | .876 | .196 | .025 | .196 |
|  | SSD | 2.817 +/- 7.883 | 1.676 | .054 | .025 | .357 |
| Cerebellum | HC | **3.145 +/- 6.441** | **2.184** | **.021\*** | **.025** | **.488** |
|  | SSD | **6.875 +/- 12.585** | **2.562** | **.009\*\*** | **.025** | **.546** |
| Cerebellum > Sham | HC | 1.985 +/- 7.617 | 1.165 | .129 | .05 | .261 |
|  | SSD | 4.058 +/- 11.499 | 1.655 | .056 | .05 | .353 |
| Active/Auditory:<br>Sham | HC | **3.699 +/- 7.323** | **2.259** | **.018\*** | **.025** | **.505** |
|  | SSD | 5.263 +/- 14.656 | 1.684 | .053 | .025 | .359 |
| Active/Auditory:<br>Cerebellum | HC | **9.181 +/- 7.356** | **5.581** | **< .001\*\*\*** | **.025** | **1.248** |
|  | SSD | **10.184 +/- 18.052** | **2.646** | **.008\*\*** | **.025** | **.564** |
| Active/Auditory:<br>Cerebellum > Sham | HC | **8.328 +/- 12.974** | **2.313** | **.016\*** | **.05** | **.517** |
|  | SSD | 4.921 +/- 20.334 | 1.135 | .135 | .05 | .242 |

Note. The TRE is defined as the difference in the percentage of detected delays between conditions with the 200ms vs. 0ms delay during preceding adaptation phases. For individual conditions, one-sample t-tests were used to assess whether the TRE was significantly greater than zero. Difference in TRE between conditions were assessed with two-samples t-tests. All t-tests were Bonferroni corrected. The corrected alpha level used for each test is displayed in the column $\alpha_{\text{corr}}$ . Significant tests are presented with bold values. \* $p < .05$ , \*\* $p < .01$ , \*\*\* $p < .001$

Regarding the influence of tDCS on temporal recalibration, according to the interaction of *stimulation* and *adaptation delay* [*F*(3, 120) = 2.800, *p* = .043, η_p_^2^ = .065] and subsequent post-hoc tests, across groups, the TRE was significantly stronger after cerebellar tDCS compared to sham stimulation [*Mean difference* = 3.071, *SD* = 8.050, *t*(41) = 2.472, *p* = .009, *d* = .381, ɑ_corr_ = .016; Sham: *Mean TRE* = 2.028, *SD* = 6.250, *t*(41) = 2.103, *p* = .021, *d* = .324, ɑ_corr_ = .025; Cerebellum: *Mean TRE* = 5.099, *SD* = 8.183, *t*(41) = 4.038, *p* < .001, *d* = .623, ɑ_corr_ = .025], but not after tDCS on the right SMA [*Mean difference* = .640, *SD* = 7.643, *t*(41) = .542, *p* = .295, *d* = .084, ɑ_corr_ = .016] or the right TPJ [*Mean difference* = .967, *SD* = 6.257, *t*(41) = 1.001, *p* = .161, *d* = .155, ɑ_corr_ = .016]. Finally, the significant four-way interaction of *stimulation, movement type, test modality*, and *adaptation delay* [*F*(3, 120) = 3.343, *p* = .022, η_p_^2^ = .077] further revealed that, across groups, the faciliatory influence of cerebellar tDCS on the TRE occurred specifically for active and auditory conditions [*Mean difference* = 5.188, *SD* = 14.052, *t*(41) = 2.393, *p* = .011, *d* = .369, ɑ_corr_ = .0125; Sham: *Mean TRE* = 4.518, *SD* = 9.580, *t*(41) = 3.057, *p* = .002, *d* = .472, ɑ_corr_ = .025; Cerebellum: *Mean TRE* = 9.707, *SD* = 12.633, *t*(41) = 4.979, *p* < .001, *d* = .768, ɑ_corr_ = .025], but was absent in passive/auditory [*Mean difference* = 3.109, *SD* = 16.246, *t*(41) = 1.240, *p* = .111, *d* = .191, ɑ_corr_ = .0125], active/visual [*Mean difference* = 1.405, *SD* = 9.725, *t*(41) = .937, *p* = .177, *d* = .145, ɑ_corr_ = .0125], and in passive/visual conditions [*Mean difference* = 2.581, *SD* = 10.710, *t*(41) = 1.562, *p* = .063, *d* = .241, ɑ_corr_ = .0125].

**Fig. 2.**
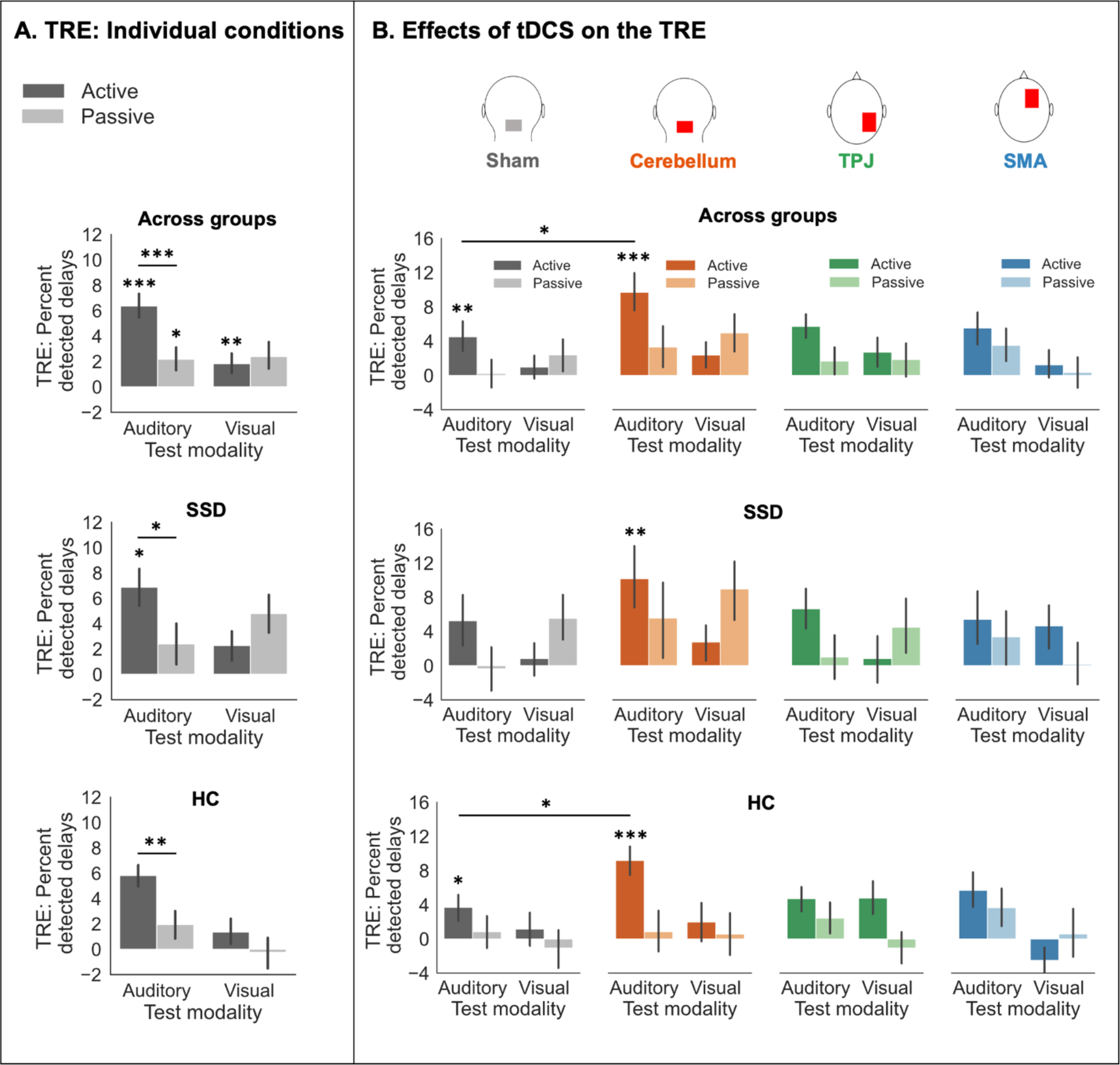
Temporal recalibration effects. **A:** The TRE, defined as the difference in the percentage of detected delays between conditions with the 200ms vs. 0ms delay during preceding adaptation phases, is displayed for each experimental condition (i.e., for both test modalities and movement types), across groups, as well as separately for both groups. In both groups, for auditory (unimodal) conditions, the TRE was significantly larger in active compared to passive movement conditions. **B:** The TRE is displayed for each of the four stimulation conditions, again across groups and separately for both groups. Across groups and for HC alone, cerebellar tDCS significantly facilitated the TRE compared to sham stimulation. For SSD, cerebellar tDCS induced a significant TRE which was absent during sham stimulation. Error bars indicate standard errors of the mean. \**p* > .05, \*\**p* < .01, \*\*\**p* < .001

## 4 Discussion

We investigated commonalities and differences in sensorimotor temporal recalibration mechanisms between HC and SSD and whether tDCS on relevant regions could facilitate recalibration effects. We found similar effects of sensorimotor temporal recalibration in both groups indicating that recalibration mechanisms may be preserved in SSD. Furthermore, the faciliatory impact of cerebellar tDCS on these effects in both groups highlights the importance of the cerebellum for recalibrating forward model predictions in response to environmental changes.

Regardless of the tDCS stimulation, both HC and SSD showed a significant TRE across conditions, and specifically so for active movements. Furthermore, no group differences in the TRE were observed depending on the movement type, test modality, or stimulation condition. Thus, our study does not provide evidence for a fundamental impairment in sensorimotor temporal recalibration in SSD, but rather suggests commonalities in recalibration mechanisms between SSD and HC. Predictable action-outcomes, i.e., outcomes in active conditions in our study, are typically associated with perceptual differences compared to externally generated sensory input.^2–4^ However, this difference is often found to be reduced in SSD which is usually considered as an indicator of impairments of the forward model in predicting the sensory outcomes of self-generated actions.^5,7–9^ In our study, overall group differences in delay detection performances between actively and passively generated stimuli appeared to have been too small or associated with too much variance to manifest in a significant interaction effect (see also supplementary material S7 for study limitations). Nevertheless, according to supplementary analyses (see supplementary material S6), SSD exhibited a reduced difference between active and passive delay detection rates for a specific test delay level, indicating that the previously reported deficit^5,7–9,35^ also weakly manifested in our data. Importantly, due to the absence of differences in temporal recalibration between the groups, our findings suggest that the aberrant processing associated with self-generated action-outcomes in SSD cannot be attributed to dysfunctions in flexibly recalibrating forward model predictions in response to changes in environmental conditions, such as varying action-outcome delays. Instead, it points to a more general failure in the prediction generation process in SSD.

Importantly, cerebellar tDCS facilitated the TRE in both groups. In HC, the TRE increased significantly with cerebellar tDCS compared to sham stimulation. In SSD, cerebellar tDCS was able to induce a significant TRE which was absent with sham stimulation. The cerebellum has frequently been suggested as the site of internal forward models.^2,4,18–21^ The adaptation of these predictions when required due to changing environmental conditions could also be associated with processes in the cerebellum.^19,28,29,48,49^ Thus, the faciliatory impact of cerebellar tDCS on the TRE suggests that the recalibration of these predictive processes in the cerebellum was amplified by the stimulation, which is in line with previous studies demonstrating a faciliatory influence of cerebellar stimulation on sensorimotor temporal recalibration mechanisms in healthy subjects.^28,29^ This is further supported by the fact that the TRE was generally larger in active than in passive conditions for both groups in our study. In both active and passive conditions, the TRE can be associated with the recalibration of the expected inter-sensory timing between the tactile sensation of the button movement and the visual or auditory outcome. A stronger TRE for active movements thus suggests that, next to inter-sensory recalibration mechanisms, the recalibration of forward model predictions additionally contributed to the TRE in this condition.^43,50^ Moreover, the fact that the faciliatory impact of cerebellar tDCS was specific to active conditions further indicates that it specifically amplified the recalibration of forward model predictions in this region.

Beyond that, the faciliatory impact of cerebellar tDCS on the TRE appeared to be specific to auditory conditions for both groups. Since the adaptation delay was always inserted between the button press and the auditory outcome, a transfer of the TRE to vision, especially for active conditions, would suggest that forward model predictions are generated and recalibrated simultaneously for sensory outcomes of different modalities.^4,51,52^ This would indicate that recalibration results in changes in the general predicted timing for sensory action-outcomes rather than in modality-specific changes.^13,43–45^ Although the adaptation procedure had an impact on temporal perception in the visual domain in our study, leading to a visual TRE across the groups, cerebellar tDCS did not affect the size of this modality-transfer effect. Furthermore, the transfer of the TRE to vision was not stronger in active than in passive conditions. Thus, these findings do not speak for the presence of supra-modal predictive mechanisms in the cerebellum. Instead, the modality-transfer can rather be explained by the supra-modal recalibration of inter-sensory matching mechanisms, assumed to be involved in both active and passive conditions, leading to changes in the expected timing between tactile, auditory, and visual outcomes.^53^

Overall, the fact that cerebellar tDCS had a similar impact on the TRE for both groups highlights the importance of cerebellum-based predictive processes, which play a vital role in the adaptation to action-outcome delays in both healthy subjects and patients with SSD. This could also imply the potential of cerebellar tDCS to facilitate related adaptive processes in SSD, which are tightly connected to the cerebellum as well and have frequently been reported to be impaired in these patients. Among them is the process of sensorimotor adaptation, i.e., the adaptation of movements in response to a discrepancy between predicted and observed sensory outcomes of these movements.^54–56^ For instance, compared to healthy subjects, patients exhibited impaired sensorimotor adaptation abilities in tasks where movements had to adapt to shifted or rotated visual feedback,^57–59^ and reduced saccade adaptation.^60,61^ These adaptation deficits in SSD may also be explained by dysfunctions in accurately building and updating internal forward models in the cerebellum to minimize the error between predicted and perceived sensory action-outcomes.^59,61^ In healthy subjects, tDCS on the cerebellum already showed the potential to improve sensorimotor adaptation performances in similar tasks.^62–64^ Initial evidence in non-clinical psychosis further demonstrated the effectiveness of cerebellar tDCS in ameliorating sensorimotor learning deficits.^65^ Since cerebellar tDCS had a faciliatory impact on sensorimotor temporal recalibration in both groups in our study, these findings emphasize the potential of this technique to also improve these related adaptive processes in SSD. Furthermore, intact sensorimotor recalibration mechanisms, which can be further amplified by cerebellar tDCS, could also represent a valuable resource of patients with SSD that could be used to train forward models to generate adequate action-outcome predictions and thus improve self-other differentiation, action-outcome monitoring, and ultimately related clinical symptoms.

In conclusion, our study points to similar sensorimotor temporal recalibration mechanisms in HC and SSD and highlights the importance of the cerebellum in both groups for this process. Our results suggest that cerebellar tDCS may constitute a promising tool for addressing deficits in related predictive or adaptive processes based on the forward model in the cerebellum, and potentially linked symptomatology in SSD.

## 5 Acknowledgements

We thank Jens Sommer for technical support and Michelle Achenbach, Trâm Đỗ, Luca Grolms, Lisa Herberstein, and Sabrina Nasri-Roudsari for assistance in data collection and patient recruitment. This research project was supported by the German Research Foundation (DFG project number: 286893149, STR1146/9-2, STR1146/15-1, CRC/TRR135, project A3) and by “The Adaptive Mind”, funded by the Excellence Program of the Hessian Ministry of Higher Education, Science, Research and Art.

## Footnotes

* Excluding the first four patients for which the adaptation delay had been set to 150ms instead of 200ms, did not yield group differences in the overall TRE either [*t*(36) = 1.508, *p* = .140, *d* = .490, two-sided], indicating that these patients did not fundamentally affect the reported temporal recalibration results.

