## Supplementary Material for "Facilitation of sensorimotor temporal recalibration mechanisms by cerebellar tDCS in patients with schizophrenia spectrum disorder and healthy subjects"

**Supplementary material** for the research article: "Facilitation of temporal recalibration mechanisms by cerebellar tDCS in patients with schizophrenia spectrum disorder and healthy subjects"

### S1. Sample characteristics

Initially, 24 patients with SSD were included in the study, but two of them dropped out before finishing all sessions due to a loss of interest in participating. The final samples of HC and SSD were matched in terms of sex and the level of education (see **Supplementary Table 1**). There were no significant age difference between the groups [independent-samples t-test: $t(40) = .343$, $p = .733$, $d = .106$].

Before the first tDCS session, patients were invited to an additional session during which their diagnosis was verified with the Structured Clinical Interview for DSM-IV (SCID).[1] Furthermore, additional clinical scales and neuropsychological tests were applied during this session (results are summarized in **Supplementary Table 1**). For HC, the same neuropsychological tests were applied during the tDCS sessions.

**Supplementary Table 1. Demographics and clinical characteristics.**

| | **SSD** (N = 22) | **HC** (N = 20) |
|---|---|---|
| ***Demographics*** | | |
| Male/Female | 11/11 | 10/10 |
| Age | 35.80 +/- 10.37 | 36.90 +/- 10.37 |
| Higher Education | 13 | 14 |
| ***Antipsychotic medication*** | | |
| None | 4 | 20 |
| FGA | 2[a] | - |
| SGA | 18 | - |
| ***Neuropsychological tests*** | | |
| Attention | | |
| d2 score | 163.23 +/- 39.43 | 175.47 +/- 43.06[b] |
| Executive functions | | |
| TMT-A (sec.) | 26.76 +/- 7.99 | 23.94 +/- 9.80 |
| TMT-B (sec.) | **72.32 +/- 22.56** | **54.51 +/- 22.90** |
| Short term memory | | |
| WAIS: FS score | 7.50 +/- 1.44 | 7.65 +/- 1.75 |
| WAIS: BS score | 6.14 +/- 1.21 | 6.60 +/- 1.43 |
| ***Clinical scales*** | | |
| SAPS score | 18.04 +/- 13.29 | |
| SANS score | 13.40 +/- 11.37 | |
| BDI score | 0.64 +/- 0.54 | |
| GAF score | 60.50 +/- 15.90 | |
| SOFAS score | 78.04 +/- 14.00 | |

*Note.* FGA = First generation antipsychotics, SGA = Second generation antipsychotics, d2: d2 test of attention,[2] TMT: Trial Making Test,[3] WAIS: Wechsler Adult Intelligence Scale,[4] FS: Forward span, BS: Backward span, SAPS:

Scale for the Assessment of Positive Symptoms,[5] SANS: Scale for the Assessment of Negative Symptoms,[6] BDI: Beck Depression Inventory,[7] GAF: Global Assessment of Functioning,[8] SOFAS: Social and Occupational Functioning Assessment Scale.[9] For continuous variables the mean +/- standard deviation is displayed. Significant differences between HC and SSD are presented with bold values [TMT-B: $t(40) = -2.536$, $p = .015$, $d = -.784$; there were no group differences for any other comparisons: all $p > .260$]. [a]Two patients were medicated with both FGA and SGA. [b]N = 19.

### S2. Electrode placement

In all active tDCS conditions, the anode was placed over the brain region to be stimulated. For stimulation of the bilateral cerebellum, the center of the anode was placed on the midline 2cm below the inion. For tDCS on the right SMA and TPJ, electrodes were positioned according to the 10-20 EEG system. For tDCS on the right SMA, the anode was placed on FC2 (10% of the distance between nasion and inion anteriorly to Cz; and 10% of the distance between the preauricular points to the right). tDCS on the right TPJ was applied by placing the anode between C4 and P4 (20% of the distance between nasion and inion posteriorly to Cz and 20% of the distance between the preauricular points to the right). During the sham stimulation session, electrodes were attached similarly as for the cerebellar tDCS condition.

### S3. Training Procedure

To ensure that button presses were correctly performed and that subjects were familiar with the task, all subjects went through a training procedure during the first tDCS session prior to the stimulation. They were trained to let their finger be moved by the button device in passive condition without applying any counter-pressure. They were also trained to perform the button presses with the correct timing, i.e., in intervals of approx. 800ms in adaptation phases and for a duration of approx. 500ms (in both adaptation and test phases). The button presses were chosen to last for 500ms to ensure that stimuli were presented before the upward movement of the button for all test delay levels (the max. delay was 417ms), since it may interfere with delay detection. Even though subjects were trained to perform the active button presses with the parameters described above, further measures were taken during the experiment to assure comparable button press parameters for active and passive movement conditions: Passive button press intervals and durations adapted to the mean of the respective preceding active conditions. Adaptation phases always terminated automatically after nine button presses in each of the two parts. If subjects completed the button presses too fast in active conditions during a part of the adaptation phase (i.e., faster than 8000ms), the jitter between the adaptation phases (when moving too fast in the first part) or the instruction text for the following test phase (when moving too fast in the second part) were extended by the remaining time. Additionally, subjects trained the test phases for each experimental condition, once with no delay and once with the maximum delay between button movement and outcome (417ms) to familiarize with the stimuli and the delays. During the training of test trials, they received feedback about the actual presence of a delay. Responses about the presence of a delay were instructed to be given as accurately, but not as fast as possible. Lastly, they went through in a 10-minute training of the experiment to further familiarize with the task.

**S4. Overview of all main and interaction effects for the main analysis**

A mixed ANOVA with the between-subjects factor *group* and the within-subjects factors *stimulation*, *test modality*, *movement type*, and *adaptation delay* was used to assess the impact of these variables on the percentage of detected delays during the test phases. Of main importance were main and interaction effects including the factor *adaptation delay* since modulations of delay detection performances were expected to occur after exposure to the delayed (200ms) vs. undelayed tone during adaptation phases. **Supplementary Table 2** provides an overview of all effects of this analysis. Please refer to the main manuscript for an interpretation and discussion of the relevant effects.

**Supplementary Table 2. Results of the ANOVA testing for differences in delay detection performances between the experimental conditions**

| Effect | Sum of Squares | df | Mean Square | F-val. | p-val. | $\eta_p^2$ |
|---|---|---|---|---|---|---|
| **Group** | 1042.879 | 1 | 1042.879 | 0.239 | 0.628 | 0.006 |
| Residuals | 174743,407 | 40 | 4368.585 | | | |
| **Stimulation** | 1889.121 | 3 | 629.707 | 3.228 | 0.025 | 0.075 |
| **Stimulation * Group** | 579.187 | 3 | 193.062 | 0.990 | 0.400 | 0.024 |
| Residuals | 23411.494 | 120 | 195.096 | | | |
| **Test modality** | 276.511 | 1 | 276.511 | 1.072 | 0.307 | 0.026 |
| **Test modality * Group** | 300.903 | 1 | 300.903 | 1.167 | 0.286 | 0.028 |
| Residuals | 10313.270 | 40 | 257.832 | | | |
| **Movement type** | 8490.601 | 1 | 8490.601 | 23.817 | < .001 | 0.373 |
| **Movement type * Group** | 15.472 | 1 | 15.472 | 0.043 | 0.836 | 0.001 |
| Residuals | 14259.492 | 40 | 356.487 | | | |
| **Adaptation delay** | 3381.799 | 1 | 3381.799 | 14.033 | < .001 | 0.260 |
| **Adaptation delay * Group** | 64.406 | 1 | 64.406 | 0.267 | 0.608 | 0.007 |
| Residuals | 9639.504 | 40 | 240.988 | | | |
| **Stimulation * Test modality** | 41.766 | 3 | 13.922 | 0.487 | 0.692 | 0.012 |
| **Stimulation * Test modality * Group** | 20.466 | 3 | 6.822 | 0.239 | 0.869 | 0.006 |
| Residuals | 3429.039 | 120 | 28.575 | | | |
| **Stimulation * Movement type** | 498.654 | 3 | 166.218 | 2.905 | 0.038 | 0.068 |
| **Stimulation * Movement type * Group** | 1.018 | 3 | 0.339 | 0.006 | 0.999 | 1.483e -4 |
| Residuals | 6865.225 | 120 | 57.210 | | | |
| **Test modality * Movement type** | 3085.947 | 1 | 3085.947 | 57.555 | < .001 | 0.590 |
| **Test modality * Movement type * Group** | 4.002 | 1 | 4.002 | 0.075 | 0.786 | 0.002 |
| Residuals | 2144.706 | 40 | 53.618 | | | |
| **Stimulation * Adaptation delay** | 442.776 | 3 | 147.592 | 2.800 | 0.043 | 0.065 |
| **Stimulation * Adaptation delay * Group** | 28.934 | 3 | 9.645 | 0.183 | 0.908 | 0.005 |

| | | | | | | |
|---|---|---|---|---|---|---|
| Residuals | 6324.609 | 120 | 52.705 | | | |
| **Test modality * Adaptation delay** | 398.065 | 1 | 398.065 | 9.229 | 0.004 | 0.187 |
| **Test modality * Adaptation delay * Group** | 8.787 | 1 | 8.787 | 0.204 | 0.654 | 0.005 |
| Residuals | 1725.286 | 40 | 43.132 | | | |
| **Movement type * Adaptation delay** | 289.127 | 1 | 289.127 | 8.762 | 0.005 | 0.180 |
| **Movement type * Adaptation delay * Group** | 74.661 | 1 | 74.661 | 2.263 | 0.140 | 0.054 |
| Residuals | 1319.897 | 40 | 32.997 | | | |
| **Stimulation * Test modality * Movement type** | 81.283 | 3 | 27.094 | 1.558 | 0.203 | 0.037 |
| **Stimulation * Test modality * Movement type * Group** | 17.785 | 3 | 5.928 | 0.341 | 0.796 | 0.008 |
| Residuals | 2086.397 | 120 | 17.387 | | | |
| **Stimulation * Test modality * Adaptation delay** | 117.841 | 3 | 39.280 | 1.425 | 0.239 | 0.034 |
| **Stimulation * Test modality * Adaptation delay * Group** | 18.754 | 3 | 6.251 | 0.227 | 0.878 | 0.006 |
| Residuals | 3307.155 | 120 | 27.560 | | | |
| **Stimulation * Movement type * Adaptation delay** | 15.263 | 3 | 5.088 | 0.131 | 0.941 | 0.003 |
| **Stimulation * Movement type * Adaptation delay * Group** | 19.708 | 3 | 6.569 | 0.169 | 0.917 | 0.004 |
| Residuals | 4656.530 | 120 | 38.804 | | | |
| **Test modality * Movement type * Adaptation delay** | 460.901 | 1 | 460.901 | 7.781 | 0.008 | 0.163 |
| **Test modality * Movement type * Adaptation delay * Group** | 33.242 | 1 | 33.242 | 0.561 | 0.458 | 0.014 |
| Residuals | 2369.495 | 40 | 59.237 | | | |
| **Stimulation * Test modality * Movement type * Adaptation delay** | 179.676 | 3 | 59.892 | 3.343 | 0.022 | 0.077 |
| **Stimulation * Test modality * Movement type * Adaptation delay * Group** | 5.751 | 3 | 1.917 | 0.107 | 0.956 | 0.003 |
| Residuals | 2149.907 | 120 | 17.916 | | | |

*Note.* $N_{HC}$ = 20, $N_{SSD}$ = 22

### S5. Stimulation side effects

After each session, subjects reported on a custom-designed questionnaire whether they perceived any side effects due to the tDCS stimulation [on a scale from one (no side effect) to five (strong side effect) for 28 items]. To test for potential differences in perceived stimulation side effects between groups and stimulation conditions, we conducted a mixed ANOVA with the between-subjects factor *group* and the within-subjects factor *stimulation*. A full overview of the results is displayed in **Supplementary Table 3**. There was a significant main effect of *group* indicating that patients with SSD (*Mean* = 1.608, *SD* = .440) reported stronger perceived side effects than HC (*Mean* = 1.229, *SD* = .281). Importantly, since there were no group differences in the TRE or in the impact of tDCS on the TRE in our study, this difference in perceived stimulation side effects should not have influenced our reported main results. There was no significant main effect of *stimulation* and no interaction of the factors *group* and *stimulation*, indicating that the amount of perceived side effects did not differ between the different stimulation conditions.

**Supplementary Table 3. Results of the ANOVA testing for differences in perceived stimulation side effects.**

| Effect | Sum of Squares | df | Mean Square | F-val. | p-val. | $\eta_p^2$ |
|---|---|---|---|---|---|---|
| **Group** | 6.030 | 1 | 6.030 | 18.459 | < .001 | .316 |
| Residuals | 13.067 | 40 | .327 | | | |
| **Stimulation** | .357 | 3 | .119 | 1.535 | .209 | .037 |
| **Stimulation * Group** | .421 | 3 | .140 | 1.810 | .149 | .043 |
| Residuals | 9.307 | 120 | .078 | | | |

*Note.* $N_{HC}$ = 20, $N_{SSD}$ = 22

### S6. Group-dependent differences in delay detection performance between active and passive conditions

The rationale of the present study was based on the established finding that patients with SSD show impairments in predicting the sensory outcomes of self-generated actions which manifests in reduced perceptual differences between actively vs. passively elicited stimuli in SSD compared to HC.[10–13] Here, we investigated whether this impairment may partly be attributed to dysfunctional sensorimotor temporal recalibration mechanisms. Thus, firstly, the question arises as to whether there was such a general impairment in predicting sensory action-outcomes in patients, meaning whether they showed a reduced difference in the processing of actively vs. passively generated stimuli in our study.

Both groups detected more delays in active compared to passive conditions. According to paired-samples t-tests, this difference was significant in HC [*Mean difference* = 6.591, *SD* = 6.561, $t(19) = 4.493$, $p < .001$, $d = 1.005$, two-sided], but failed to reach significance for SSD [*Mean difference* = 3.635, *SD* = 9.135, $t(21) = 1.866$, $p = .076$, $d = .398$, two-sided]. Nonetheless, this group difference appeared to be not strong enough or the variance within the groups might have been too large to lead to a significant *group* x *movement type* interaction in the ANOVA reported in the main manuscript.

However, since it may be reasonable to assume that the active-passive difference in delay detection occurs for certain delay levels only, we also conducted a generalized estimating equations (GEE) analysis using IBM SPSS Statistics (Version 27.0) with the active-passive difference in the percentage of detected delays as dependent variable, which was computed separately for each of the six delay levels used during the test phases. An AR (1) working correlation structure and robust (sandwich) covariance estimators were used for the regression coefficients. The factors, *stimulation* (cerebellum, TPJ, SMA, sham), *test modality* (auditory, visual), *test delay* (0, 83, 167, 250, 333, 417ms), and *group* (HC, SSD) were included in a full factorial model testing for all main and interaction effects. The active-passive difference was modeled with a linear link function. Importantly, this analysis revealed a significant interaction of *group* and *test delay* [Wald Chi-Square (df = 5) = 13.332, $p$ = .020]. According to post-hoc tests, the active-passive difference was significantly stronger in HC than in SSD for test stimuli delayed by 250ms [mean difference = 11.512, standard error = 3.928, df = 1, $p$ = .003].

Hence, for an individual medium-sized delay level, patients showed reduced differences in the perception of actively vs. passively elicited stimuli. Importantly, the medium-sized delay levels are the ones for which most prominent active-passive differences can be assumed due to floor or ceiling effects at very small or large delays, respectively. Thus, to a given extent, there are indications for the aberrant processing of actively generated action-outcomes in SSD in in our study, but we cannot provide evidence for the attribution of this impairment to dysfunctional temporal recalibration mechanisms.

**S7. Limitations**

The results of our study do not provide evidence for impaired sensorimotor temporal recalibration mechanisms in patients with SSD, as there were no significant differences compared to the HC group. This could indicate that sensorimotor recalibration abilities may be a useful resource of patients with SSD that could be exploited to train predictive mechanisms based on the forward model and to thereby improve self-other differentiation and action-outcome monitoring. Importantly, however, the absence of group differences in recalibration does not necessarily speak against the existence of an impairment in patients; it is also conceivable that certain characteristics of our study have masked differences between the groups.

Firstly, previous studies on potentially related adaptive processes, namely on sensorimotor adaptation, could show that patients were able to adapt their movements to the introduced action feedback perturbations, but they adapted slower[14] and the adaptation process was associated with more errors[15] compared to HC. Our study design did not allow for the investigation of the time course of temporal recalibration effects. Adaptation and test phases were blocked, which only allowed us to assess the impact of the adaptation delay on perception once at the end of an adaptation phase. Thus, future study designs should consider that the time course of adaptation could provide important information regarding potential impairments in patients.

Secondly, the majority of patients in our sample were under antipsychotic medication at the time of the study. This could have compensated for potentially existing deficits, since antipsychotics are known to particularly target positive symptoms, such as hallucinations and delusions, which are believed to be associated with the dysfunctions in predictive mechanisms of the forward model investigated here.[10–13,16,17]

Thirdly, the variance in temporal recalibration and stimulation effects appeared to be high in our study, particularly in the patient group. While this may partly be explained by the relatively low sample sizes, it could also suggest that there are individual differences in whether patients exhibit dysfunctions in sensorimotor temporal recalibration mechanisms. Likewise, there could be individual differences in the effectiveness of tDCS on enhancing recalibration effects. Hence, it would be useful to determine under which conditions patients may exhibit dysfunctions in temporal recalibration or profit from tDCS. A relevant factor could, for instance, be the presence the above-mentioned symptoms related to the presumed deficits in predictive mechanisms. Future studies with larger sample sizes could specifically investigate the determining factors for the occurrence of a potential deficit in this process and the effectiveness of tDCS on facilitating the underlying predictive mechanisms. Furthermore, variability in stimulation-dependent effects may also arise due to individual differences in subjects´ cerebellar anatomy and connectivity pattern with other brain regions.[18] Thus, future studies could apply individually adjusted stimulation protocols according to the subjects´ individual anatomy.
